# Transfer learning enables discovery of sub-micromolar antibacterials for ESKAPE pathogens from ultra-large chemical spaces

**DOI:** 10.1101/2025.08.07.669197

**Authors:** Miguel García-Ortegón, Srijit Seal, Emily Geddes, Jenny L Littler, Collette S Guy, Jonathan Whiteside, Carl Rasmussen, Andreas Bender, Sergio Bacallado

## Abstract

The rise of antimicrobial resistance, especially among gram-negative ESKAPE pathogens, presents an urgent global health threat. However, the discovery of new antibiotics is hampered by sparse publicly available antibacterial data, complex bacterial defenses, and weak economic incentives. Here, we introduce a transfer learning framework using deep graph neural networks (DGNNs) to identify antibacterials from ultra-large chemical libraries. DGNNs were first pre-trained on large molecular datasets of protein-ligand simulations, binding affinities, and physicochemical properties to learn generalizable chemical features, and then fine-tuned on limited antibacterial screening data. Compared to classical methods, transfer learning significantly improved enrichment factors and predictive performance in cross-dataset benchmarks. Applying this strategy to the ChemDiv and Enamine libraries, we virtually screened over a billion compounds and prioritized 156 candidates. Experimental testing against *Escherichia coli* revealed that 54% of compounds exhibited antibacterial activity (MIC ≤64 µg/ml), with several demonstrating sub-micromolar potency and broad-spectrum efficacy against gram-positive and gram-negative pathogens, including three ESKAPE species. Of 18 broad-spectrum candidates, 15 showed minimal cytotoxicity and no hemolytic activity. These results validate our approach for navigating underexplored chemical space and identifying potent, low-toxicity compounds with antibiotic activity. We release open-source models and a scalable workflow to accelerate antibacterial discovery in the face of data scarcity.

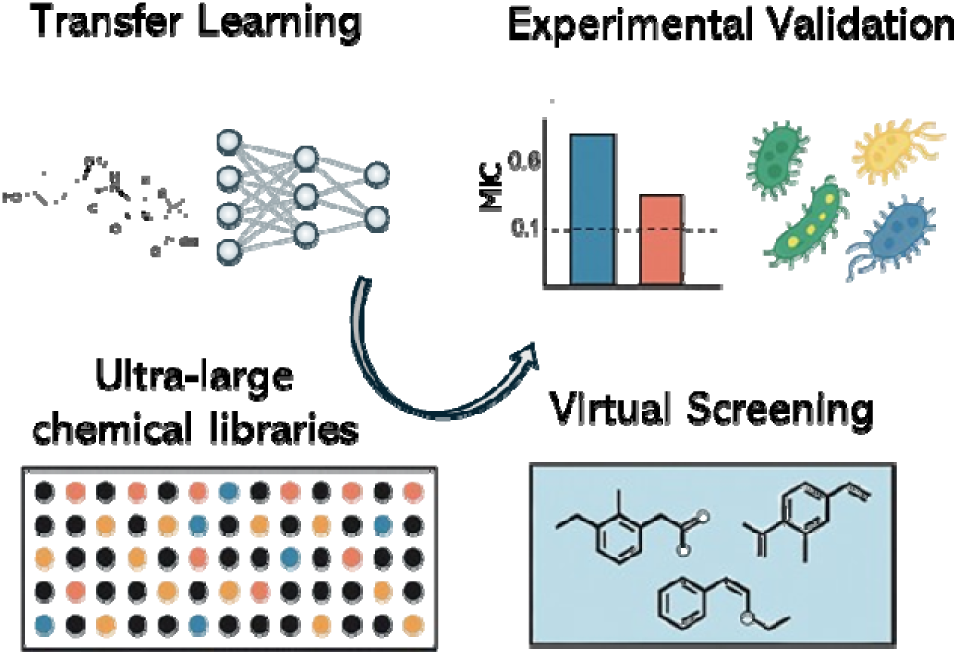

## Introduction

The global rise of multidrug-resistant pathogens is a profound crisis, with 1.27 million deaths directly attributed to resistant bacteria in 2019^1^ and a projected 10 million deaths per year by 2050^2^. Infections caused by gram-negative bacteria are particularly challenging to treat due to the limited diversity of antibiotics, and the FDA has not approved any new class of antibiotics for treating gram-negative infections in over 50 years ^3^. One significant challenge in antibiotic discovery is that gram-negative bacteria possess a highly impermeable outer membrane and active efflux pumps, which work together to prevent intracellular accumulation of many compounds. This dual defense mechanism severely limits the effectiveness of traditional target-based drug design strategies, as potential antibiotics often fail to reach their intended intracellular targets in sufficient concentrations to exert their effects.^4,5^ Further, the incentives for antibiotic development are poor^6,7^, and there is an inherent bias against complex chemistries and natural products that are typically not considered drug-like^7^. Therefore, cost-effective and efficient methods for identifying novel chemical matter with whole-cell activity against Gram-negative bacteria are highly desirable.

Machine learning models are increasingly being utilized to predict the bioactivity and toxicity of compounds.^8–10^ Deep neural networks (DNNs) have recently arisen as a powerful tool for virtual screening to discover structurally distinct compounds with antibacterial activity. Notable examples are the work by Stokes et al., who used ensembles of deep graph neural networks (DGNNs) to discover halicin, an antibiotic with a new mechanism of action^11^, and Wong et al., who used similar models to find a novel structural class of antibiotics^12^. DNNs are appealing because, as deep overparameterized models, they can learn rich, continuous representations of various data types, including discrete ones. This ability makes them highly flexible and enables high predictive performance in several application domains, including image recognition^13^, image generation^14^, and natural language processing^15^, among others. However, unlike these fields, where abundant public data is available (e.g., images, videos, and text), the availability of labeled positive examples in antibiotic datasets (i.e., compounds with antibiotic properties) remains exceedingly limited. This scarcity significantly constrains supervised training, particularly for high-capacity models. For example, the crowdsourced dataset COADD^16^ for *E. coli* ATCC 25922 contains just 159 active compounds (considering an 80% growth inhibition threshold, Table 1), and many of these are structural analogs, with approximately 15 unique antibiotic classes represented. This limited data set hinders the practical application of DNNs, which generally require large amounts of data to achieve high predictive accuracy. Further, these models suffer from overfitting in the low-data regime.

**Table 1.** Training and pre-training datasets. The number of scaffolds was determined using Bemis-Murcko (BM) scaffolds.

| Usage | Dataset | Description | Number of Targets | Number of Compounds | Number of Scaffolds | Number of Actives in Antibacterial Datasets | Completeness |
| --- | --- | --- | --- | --- | --- | --- | --- |
| Pre-training | RDKit | In-silico physicochemical properties | 208 | 877k | 255k | n/a | 100% |
| Pre-training | ExCAPE | Binary labels against human targets | 1,332 | 877k | 255k | n/a | 5.90% |
| Pre-training | DOCKSTRING | Docking scores against human targets | 58 | 260k | 100k | n/a | 100% |
| Training, fine-tuning | Stokes | Bacterial growth | 1 | 2,220 | 1,066 | 118 | 100% |
| Training, fine-tuning | COADD | Bacterial inhibition | 1 | 81,225 | 23,843 | 159 | 100% |

Transfer learning is a model training strategy that aims to increase the performance of machine learning models in the absence of sufficient training data and has proved effective in molecular property prediction tasks with graph neural networks.^17^ In this approach, models are not trained on the final task of interest from scratch, but rather, they are trained in two stages. First, they are pre-trained on separate tasks for which large amounts of data are available. Ideally, these tasks should be highly related to the final task of interest; however, this is not required^18^. In the second step, the parameters learned during pre-training are adapted to the final task of interest, for which training data is scarce, in what is known as fine-tuning. Crucially, fine-tuning involves minor modifications of the pre-trained parameters to avoid overfitting to the limited dataset of the task of interest. This can be achieved by setting a low learning rate or a smaller number of epochs. In this way, transfer learning attempts to learn general representations during pre-training and subsequently adapts them to maximize performance on downstream, specific tasks.^19^

Recent advances in transfer learning have leveraged knowledge gained from large-scale, often general-purpose datasets to improve performance on specific downstream tasks. Li et al. demonstrated the potential of MolPMoFiT, a fine-tuned language model for molecular activity prediction, marking a pivotal step toward next-generation QSAR modeling through inductive transfer learning.^20^ Similarly, King-Smith et al. introduced a foundational model for chemistry, showing how large pre-trained architectures can be adapted across a wide spectrum of chemical tasks.^21^ In the domain of chemical reactivity, Keto et al. leveraged chemically aware pre-training to improve reaction prediction performance^22^, while Noto et al. successfully transferred learned representations across different photocatalytic reaction classes^23^. Overall, this shows that domain-specific transfer learning strategies can be used to improve models, and in this work we aimed to use transfer learning to enhance virtual screens for sub-micromolar inhibitors of ESKAPE pathogens, whereafter the model predictions were experimentally validated.

Here, we take a transfer-learning approach to train an ensemble of DGNNs for virtual screening in order to identify compounds with activity against the gram-negative bacterial species *Escherichia coli*. Our training workflow (Figure 1a) comprised two stages: during pre-training, models were optimized to learn general molecular labels that were not specific to bacteria, such as docking scores, binding affinity against and physicochemical properties. During fine-tuning, models were optimized on small public antibacterial datasets measured on *E. coli*. Our virtual screening protocol (Figure 1b) prioritized the top predictions by the trained DGNN ensemble while maximizing the diversity of the final subset. First, we selected the highest-ranking compounds from two large commercial libraries, Chemdiv and Enamine, using clustering based on fingerprints and grouping based on antibiotic class or functional group to increase chemical diversity. Second, we validated our protocol by testing the antibacterial activity of the hit compounds against *E. coli*, finding high enrichment of actives. Finally, we also tested against a panel of gram-positive and gram-negative species, including multidrug-resistant strains of ESKAPE pathogens. Our protocol identified several compounds that were structurally novel, nontoxic and demonstrated broad-spectrum activity.

**Figure 1.**
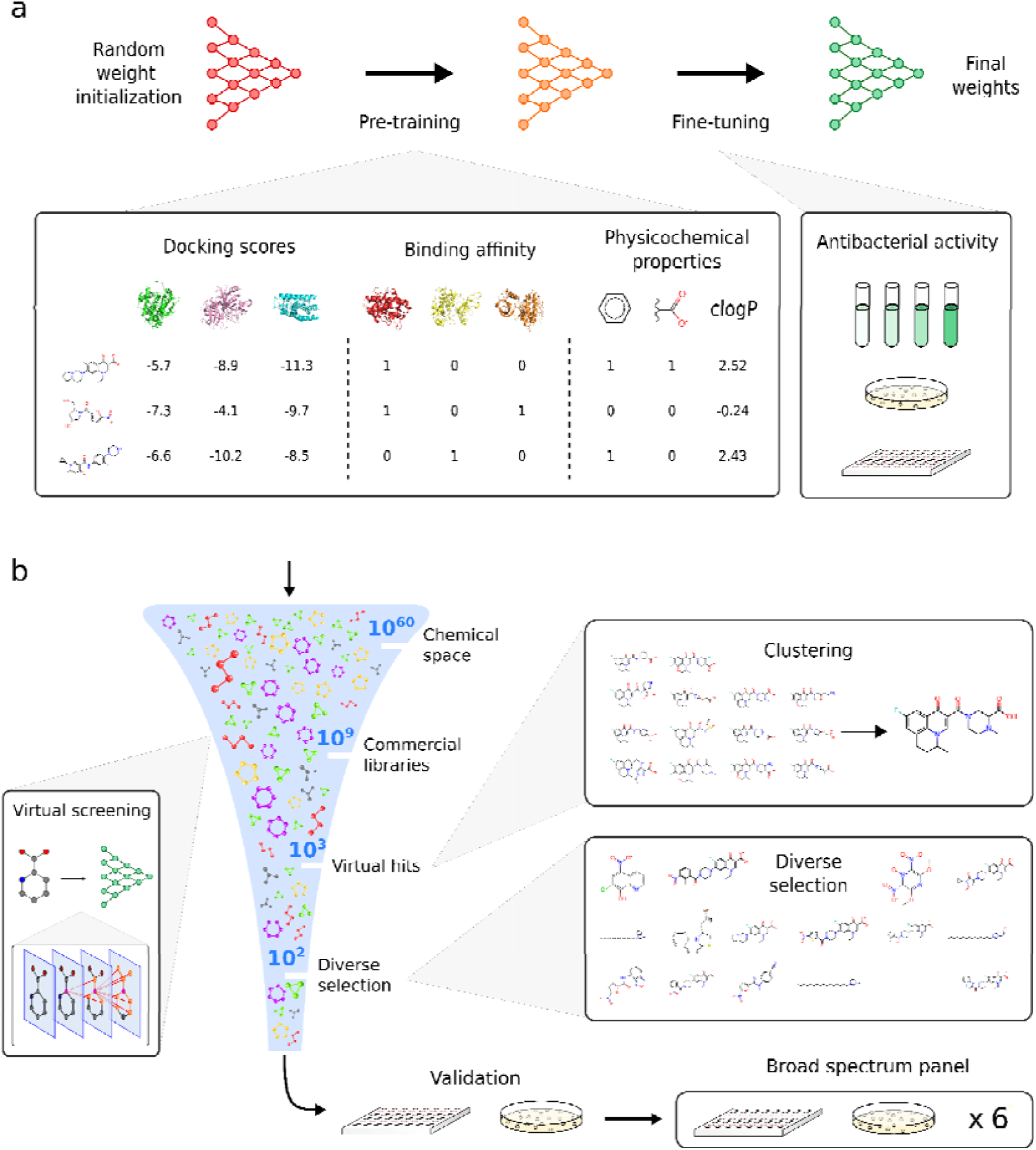
Workflows for transfer learning (a) and virtual screening of large chemical libraries (b). Before fine-tuning antibacterial data, graph neural networks were pre-trained on various molecular data, including docking scores, binding affinity, and physicochemical properties. During the virtual screening, molecules from large libraries were prioritized according to their predicted inhibition, and clustering was used to achieve a diverse candidate selection to increase the chances of finding structurally unique hits. Candidates were first validated by testing antibacterial activity against a single bacterial strain, and active candidates were further tested against a panel of gram-positive and gram-negative bacteria.

## Methods

### Datasets

We used molecular property datasets for pre-training and specific antibacterial datasets for fine-tuning or training (Table 1). Due to limited public data from antibacterial experiments, our pre-training datasets were not highly related to antibacterial activity. Still, where pre-training datasets are sufficiently numerous and diverse, both in terms of molecular structure and task labels, they are known to promote the learning of general molecular representations that can benefit fine-tuning on downstream bacterial tasks.^24^

For pretraining, we curated a dataset of RDKit molecular descriptors (calculated using RDKit v2020.09.5), ExCAPE binding affinity annotations (v2 2019), and DOCKSTRING docking scores^25,26^. These datasets, docking scores, binding affinities, and physicochemical properties, were selected as pretraining targets because they represent general, transferable molecular features relevant across many bioactivity domains and are well-represented in public datasets. These properties have been shown to be useful in transfer learning using chemically-informed feature space prior to fine-tuning on scarce data.^27,28^ We used the ExCAPE data aggregating affinity assays from PubChem and ChEMBL^29^, particularly binary affinity labels against mammalian proteins from physical experiments, although it is highly sparse. All molecular structures were stored as SMILES strings. We used DOCKSTRING, which contains AutoDock Vina docking scores of a subset of 260k ExCAPE molecules against human protein targets.^25^ For all datasets above, stereochemical information was discarded for consistency since not all datasets included it. During hyperparameter optimization experiments, SMILES were standardized using the same pipeline as DOCKSTRING^25^, which included canonicalization with RDKit^30^, discarding SMILES with unnatural charges (formal charges on atoms other than N or O), explicit hydrogens, radicals or multiple molecular fragments, and finally, protonating SMILES at pH 7.4 with OpenBabel^31^. We calculated RDKit chemical descriptors^30^ for all molecules in the ExCAPE dataset. Descriptors, as implemented in rdkit.Chem.Descriptors^30^ module included physicochemical and topological properties such as molecular weight, number of valence electrons, maximum and minimum partial charges, Bertz complexity index, logP, and number of rings (Figure 1a). Finally, this resulted in a dataset, RED, a concatenation of RDKit, ExCAPE, and DOCKSTRING features for the 260k overlapping molecules from DOCKSTRING.

For fine-tuning and training, we used the antibacterial datasets by Stokes et al. (hereafter referred to as the ‘Stokes dataset’) and the Community for Open Antimicrobial Drug Discovery (COADD).^7,16^ Both datasets were generated in wild-type *E. coli* strains, with Stokes et al. using *E. coli* BW25113 and COADD using *E. coli* ATCC 25922. We focused on *E. coli* ATCC 25922 for this study. Antibacterial activity can be expressed in potency using concentration units or growth inhibition using values between 0 (no inhibition) and 1 (full inhibition). Growth inhibition is standard in high-throughput experiments exposing bacteria to a fixed, constant compound. Stokes indicated results regarding final growth achieved rather than final inhibition (i.e., in the raw Stokes dataset, zero would indicate the highest level of inhibition and one the lowest), while COADD expressed values as percentages. We processed Stokes and COADD values to make their notation consistent (Figure 2). Occasionally, values outside the usual 0-1 range were observed: negative values indicated that the compound promoted bacterial growth rather than halting it, whereas values slightly above one could result from experimental variability or error. For classification, inhibition values were binarized using an activity threshold of 0.8 (active if inhibition > 0.8), consistent with the one previously employed by Stokes et al. All datasets are released via https://github.com/mgarort/dockbiotic/tree/main/data/ for public use.

**Figure 2.**
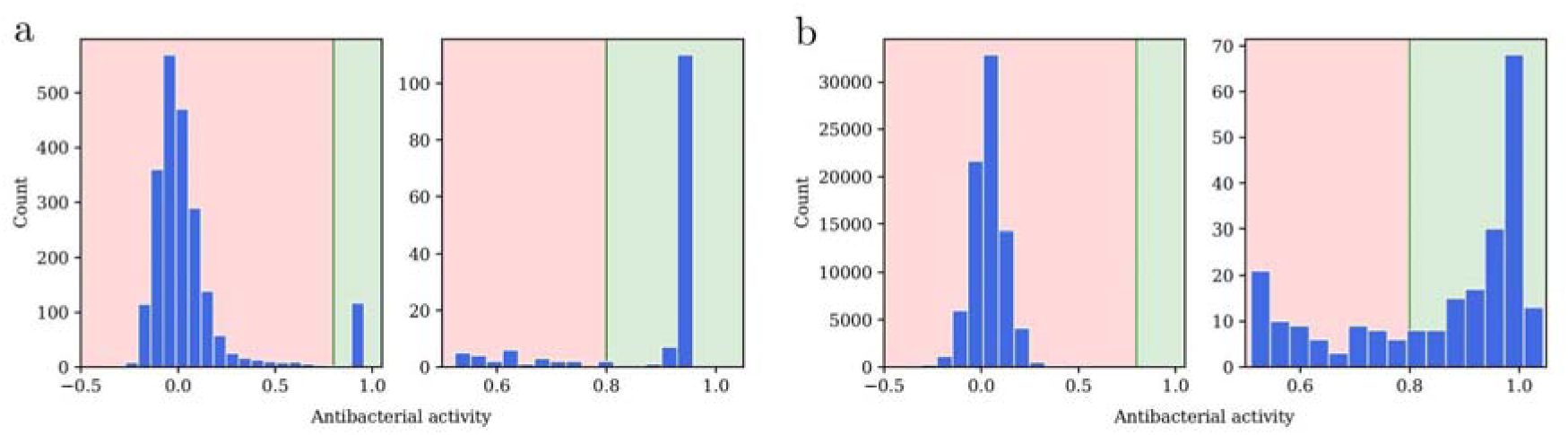
Measured bacterial inhibition on *E. coli* from the (a) Stokes and (b) COADD dataset, processed here to make the notation consistent. A value of 0 indicated no inhibition or full growth, whereas 1 indicated full inhibition or no growth. Note that measurements outside the [0,1] range were possible: values below 0 suggested that the molecule promoted bacterial growth, while values slightly above 1 could result from experimental variability or error. Count refers to the number of compounds.

### Model Architecture

We benchmarked two models: XGBoost, a tree-based ensemble on molecular fingerprints, and AttentiveFP, a deep graph neural network (DGNN) on molecular graphs.

XGBoost (eXtreme Gradient Boosting) is a popular algorithm for training tree ensembles for regression or classification. The key idea of tree ensembling is to combine several weak learners (simple models slightly better than random guessing) into a single strong learner (a model with high predictive performance) that takes a majority vote. Boosting means that the ensemble is trained by adding one tree at a time, and new trees are built, giving more weight to the training data points that are incorrectly classified. Thus, complex examples become more influential as training progresses. We employed the original XGBoost implementation^32^, taking binary Morgan fingerprints of length 2048 or 4096 as inputs.^30,33,34^

Attentive FP^35^, a DGNN, generates a vector representation of a molecule in two steps. First, it produces an embedding of each atom in an iterative procedure of message passing with attention. Each atom embedding gets updated at each iteration through a transformation that takes in that atom’s current embedding and its neighbors’ current embeddings (message passing). The influence of all neighbors in the transformation is not equal, but rather, they are weighted by attention coefficients. Second, it produces an embedding of the entire molecule in another iterative procedure with attention. An initial embedding of the molecule is produced by summing all the atoms’ embeddings. Then, for several iterations, the molecular embedding gets updated through a transformation that takes in each atom’s current molecular embedding and the current molecular embedding. Attention coefficients weigh the influence of each atom in the transformation. Finally, once the molecule embedding has been generated, a prediction can be performed with a final linear layer. We used the DeepChem^36^ implementation of Attentive FP. The input for Attentive FP was DeepChem graph representations of type *MolGraphConvFeaturizer* using edges, which incorporates bond features by concatenating one-hot vectors for bond type (single, double, triple, or aromatic), same ring membership, conjugation status, and stereo configuration. These edge features modulate the attention-weighted aggregation of neighbor atom embeddings during message passing, allowing the model to distinguish between different bond types when computing molecular representations.

All models used a batch size of 64 and were trained with Adam optimization.^37^ When training with transfer learning, we used a learning rate of 10^−3^ for pre-training and a learning rate of 10^−4^ for fine-tuning. All model weights are unfrozen during fine-tuning, allowing the full network to adapt to the antibacterial task. This allows the final linear prediction layer and upper attention mechanisms to likely undergo the most significant changes during fine-tuning, while early atom embedding layers change less. Because lower layers capture fundamental molecular features that transfer well across tasks, higher layers need to adapt for specific prediction targets. The 10x learning rate reduction preserves useful pre-trained chemical representations while allowing task-specific adaptation, preventing catastrophic forgetting while enabling efficient domain transfer. When trained without transfer learning, training was consistent with the fine-tuning phase of transfer learning to facilitate comparisons. Therefore, models not pre-trained were trained with a learning rate of 10^−4^ throughout.

### Loss functions

For the losses used during the pre-training phase, we employed MSE for the regression of absolute values (DOCKSTRING and RDKit) and cross-entropy to classify binary values (ExCAPE). ExCAPE was highly sparse; we ignored missing ExCAPE values when calculating the loss.

Inhibition values in the processed COADD and Stokes datasets were real numbers ranging between 0 and 1. The prediction of these values could be framed as regression or binary classification after binarizing according to some threshold. However, both of these settings are simplifications and carry potential disadvantages. For example, consider a molecule with a measured inhibition of 0.7. Predictions of 0.4 and 1.0 would be assigned the same quantitative error by a regression loss even though 1.0 is qualitatively very different because it indicates perfect inhibition. Similarly, consider the binary classification setting and a binarization threshold of 0.8. Molecules with measured inhibition of 0.1 or 0.7 would both be binarized as inactive, even though these values indicate very different levels of growth. For these reasons, in addition to losses for regression (MSE) and binary classification (cross-entropy), we defined a custom loss function to predict inhibition values between 0 and 1, which we call the inhibition loss. The inhibition loss combined regression and classification properties by using a custom sigmoid-based squashing function centered at a threshold (e.g., 0.8) to approximate a binary classification loss. It penalized false positives and false negatives differently using adjustable scaling factors c^+^ and c^−^, ensuring flexibility in prioritizing errors (for further details, see Supplementary Information S1).

### Model benchmarking

We evaluated different prediction model classes, losses, and transfer learning protocols to find an optimal high-performing model for virtual screening to discover antibiotics. Specifically, we benchmarked (1) XGBoost and AttentiveFP models, (2) mean squared error (MSE) for regression, cross-entropy for binary classification, and the custom inhibition loss for inhibition values between 0 and 1, and (3) training from scratch without pre-training or with pre-training on the RDKit, ExCAPE, DOCKSTRING, or a combination of the three datasets. Further, we benchmarked the amount of pre-training epochs (between 1 and 20) and fine-tuning epochs (100 and 1000). A fixed number of ten pre-training and ten fine-tuning epochs was used to benchmark other parameters.

In virtual screening, molecules from an extensive library are ranked computationally, and the top subset is selected and tested. In our benchmarking experiments, we trained all models on the Stokes dataset and tested them on a reduced version of COADD, selecting the top-ranked compounds from COADD to emulate a virtual screening workflow. COADD was reserved for testing because it was more extensive and diverse (with respect to the number of scaffolds, see Table 1) than Stokes, having been aggregated from compounds suggested by numerous independent research groups over several years. Therefore, it is expected to better represent the screening of an extensive chemical library. We removed all compounds from COADD that were analogs or structurally similar to those in Stokes, as determined by Tanimoto similarity computed on Morgan fingerprints and RDKit path fingerprints. Compounds with a Tanimoto similarity higher than 0.9 on any of the two fingerprint types were removed.

Model performance for virtual screening was quantified with the enrichment factor (EF)which is defined as the ratio of actives in the top selected subset over the ratio of actives in the initial library:

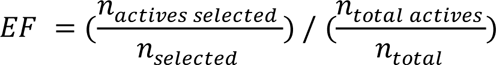

We computed the EF on the top 200 compounds from COADD. Specifically, we computed the average EF over three random repetitions with different initializations for each hyperparameter combination.

### Training final models

Once the optimal model and transfer-learning hyperparameters had been selected, the final models for virtual screening were fine-tuned on both Stokes and COADD so that they could learn from all available data. COADD, with an active rate of 0.002, was substantially more imbalanced than Stokes, which had an active rate of 0.05. To maintain a similar active rate while training the final models, as during hyperparameter optimization, we upsampled the active molecules of COADD 100 times.

We trained an ensemble of 6 models using the two best transfer-learning hyperparameters (3 models with different random seeds for each hyperparameter combination). We trained two ensembles of 6 models: one with fully standardized SMILES for ChemDiv and one with minimally standardized SMILES for Enamine since Enamine SMILES were too numerous (5.52b) to undergo complete standardization. Here, fully standardizing SMILES involved removing isomeric information, protonation, and validation for a consistent, chemically valid representation while minimal standardization of SMILES only removed isomeric information and canonicalized the structure without additional processing. The latter is used for larger datasets (like Enamine) to save on computational resources.

### Virtual screening

#### Screening libraries

We screened the commercial libraries ChemDiv^38,39^ and Enamine REAL (REadily AccessibLe)^40^. The former is an in-stock chemical collection with millions of compounds. In contrast, the latter is a combinatorial list of billions of molecules predicted to be synthesizable from initial reagents and chemical reactions (Table 2). Therefore, not every compound ordered from Enamine can eventually be acquired; their self-reported synthesis success rate is 80%.^41^

**Table 2:** Description of Screening libraries used in this study.

| Library | Version | # Compounds | Number of Bemis-Murcko Scaffolds |
| --- | --- | --- | --- |
| ChemDiv | 1566731 | 1.57M | 277k |
| Enamine | 2022q1-2 | 5.52B | 2.94B (estimated) |

For ChemDiv (1.57 m compounds) SMILES were fully standardized using the same protocol from DOCKSTRING. Enamine’s (5.52b) large size precluded charge checking and protonation, so SMILES in pre-training, training, and screening underwent a minimal standardization protocol of canonicalization with RDKit.

XGBoost employed a fingerprint molecular representation computed with RDKit, and AttentiveFP used a graph representation by DeepChem.^36^ Molecular fingerprints were also used to calculate Tanimoto similarity to assess closeness to the training set. We computed binary Morgan fingerprints (2048 bits) and RDKit fingerprints of path length six and length 2048, calculated the Tanimoto similarity on each fingerprint type, and used the highest of the two as the similarity score.

We reported the number of Bemis-Murcko (BM) scaffolds in each dataset using RDKit (Table 2). The Enamine dataset was too large to decompose every molecule, so we estimated the number of BM scaffolds on a random sample of 2 million compounds.

#### Selection Protocol

For our screening protocol, we first ranked the compounds in ChemDiv and Enamine by the inhibition activity predicted by the final ensemble of DGNN models. We selected molecules with mean predicted inhibition above 0.5. Second, we removed candidates with Tanimoto similarity (computed on Morgan fingerprints and RDKit path fingerprints) higher than 0.8 to any molecule in the COADD and Stokes training sets. By doing this, we attempted to avoid selecting compounds too structurally similar to known antibiotics. Third, many of the top-ranking compounds in Enamine were potentially redundant, with many structurally similar molecules. To increase the diversity and reduce the number of molecules selected, we clustered our selection with DBSCAN^42^, using RDKit path fingerprints of length 2048 and ε= 4.5, and kept the five top-ranking compounds within each cluster. Finally, we grouped the selected molecules by antibiotic class or functional group and selected top-ranking molecules within each group. Compound Selection Protocol. We employed DBSCAN clustering for compound selection because it determines the optimal number of clusters and demonstrates robustness to noise, making it particularly suitable for working with structurally diverse, high-dimensional data, though we acknowledge that alternative methods like K-means or agglomerative clustering could be used since the clustering approach is secondary to the transfer learning methodology itself.

#### Visualization of learned chemical space

One of the advantages of neural models is that they can learn abstract continuous representations of discrete data types such as molecules. This ability has been previously studied to enable the exploration of chemical space with generative models such as molecular VAEs.^43^ Purely supervised methods can also benefit from representation learning because clustering similar compounds within nearby regions of representation space may facilitate making predictions. AttentiveFP, the DGNN model in our ensemble, calculates predictions by first generating embeddings for each atom and later producing an embedding for the whole molecule by aggregating atomic representations. To better understand whether the AttentiveFP models trained for virtual screening were capturing antibiotic class-specific features, we performed dimensionality reduction of the molecular representations learned by AttentiveFP. Using one of the models in the ensemble, we produced AttentiveFP embeddings (200-dimensional) of training molecules (all Stokes molecules and a random sample of 50k COADD molecules) and screening molecules (a random sample of 50k ChemDiv molecules). Then, we computed 2-dimensional embeddings with UMAP^44^, using default parameters, and visualized them in a scatter plot.

### Experimental validation

**Bacterial strains:** All tested compounds were evaluated for antibacterial activity against the wild-type *E. coli* strain 25922 from the National Collection of Type Cultures (NCTC). A subset of compounds was assessed in a panel of strains, including a uropathogenic strain of *E. coli* (ECU) 13400, *P. mirabilis* (PM) 432002, *K. pneumoniae* (KP) 13442, *A. baumannii* (AB) 19606 and *S. aureus* (SA) 29213. The strains EC 13400, AB 19606, KP 13442, and SA 29213 were obtained from the NCTC, and the strain PM 432002 was obtained from the American Type Culture Collection (ATCC).

#### MIC and MBC determination protocols

The antibacterial activity of the candidate compounds was evaluated using minimum inhibitory concentration (MIC) and minimum bactericidal concentration (MBC). MIC indicates the lowest compound concentration that halts growth, while MBC refers to the lowest compound concentration that kills all bacterial cells. The Warwick Antimicrobial Screening Facility conducted these experiments using CLSI and EUCAST guidelines on antimicrobial susceptibility testing. Bacteria were cultured in cation-adjusted Mueller Hinton broth at 37 °C. For MBC experiments, bacteria were plated on cation-adjusted Mueller Hinton agar plates. Two antibiotic controls, ampicillin and ciprofloxacin, were used in this experiment. The growth and no growth control were media and bacteria without antibiotics and media alone, respectively.

In brief, a 2-fold serial dilution of each molecule in DMSO was prepared from 256 µg/ml down to 0.000122 µg/ml across two 96-well plates. A bacterial culture was prepared following the MacFarland 0.5 standard and added to each well. Plates were incubated for 18 hours at 37 °C without shaking. The MIC was the concentration of the last well with complete inhibition (i.e., a completely clear well). For the MBC, 10 µl of each well was pipetted onto agar plates and further incubated for 24 hours at 37 °C without shaking. The formation of colonies was observed, and the concentration of the last well from which no colonies were formed was taken as the MBC.

### Cytotoxicity assays

HepG2 cells were cultured in DMEM supplemented with 10% fetal bovine serum (FBS) and seeded at 4×10^4^ cells per well in a 96-well tissue culture plate. After incubation at 37°C with 5% CO_2_ for 24h, the media was aspirated off and replaced with 98 μL of fresh media. Compounds were serially diluted 2-fold in DMSO to give stock concentrations ranging from 3.2 mg/mL to 200 μg/mL, and then 2 μL of compound stock was added to the plates, resulting in a final DMSO concentration of 2%. Final compound concentrations ranged from 64 μg/mL to 4 μg/mL. After a 24-hour incubation, the cytotoxicity of compounds was evaluated using the CytoTox-GloTM assay (Promega) following the manufacturers’ instructions. Luminescence was measured using the GloMax® plate reader (Promega). Cells receiving media + 2% DMSO were used as a negative control, and cells receiving media + 2% Tritonx100 were used as the positive control for maximum cell cytotoxicity.

### Hemolysis assay

Defibrinated equine blood was centrifuged (1000×g, 10 min), and the supernatant was removed. Pelleted erythrocytes were washed thrice with PBS and resuspended to a 5% erythrocyte concentration in PBS. Compounds were serially diluted 2-fold in PBS to give a range of concentrations from 256-0.25 μg/mL, 100 μL total volume. Controls of PBS, equivalent DMSO concentrations, and Triton X-100 were also added to the plates. The resuspended erythrocytes (100 μL) were added to each well and incubated without shaking (37°C, 1 hr). Plates were then centrifuged (1000xg, 10 min), and 50 μL of supernatant was removed and added to 50 μL of PBS in a fresh plate. The absorbance was then measured at 550 nm, and the hemolytic concentration for each compound was determined, where the lowest concentration deemed to cause >10% cell lysis was judged to be the hemolytic concentration (PBS control used to represent 0% lysis and Triton X-100 used at 100% lysis). Each measurement was performed in triplicate.

## Results and Discussion

### Few active compounds are present in antibacterial dataset

Public datasets contain few compounds with high antibacterial activity and these datasets are often quite imbalanced (especially for gram-negative bacteria, where for every one active compound reported, there are hundreds of inactive compounds). For example, the database AntibioticDB records only 2939 molecules with antibacterial activity.^45^ This makes the datasets insufficient for training neural networks which need hundreds of thousands of datapoints. To increase the learning efficiency of our DGNN models, we adopted a transfer approach, using datasets with hundreds of thousands of molecules and hundreds of millions of labels for pre-training. Our pre-training labels were not directly related to the antibacterial activity; instead, they represented general physicochemical properties (RDKit), affinity binding labels against mammalian proteins (ExCAPE,) or docking scores against human protein targets (DOCKSTRING) (Table 1). Thus, pre-training aimed to learn general molecular representations that could be fine-tuned on antibacterial activity data later. For training or fine-tuning, we processed and homogenized the COADD and Stokes datasets, which represented inhibition as values between 0 (no inhibition) and 1 (complete inhibition) (Figure 2). Most compounds in these datasets were inactive, with only 118 molecules in Stokes (screened at 50 μM) and 159 molecules in COADD eliciting inhibition above the 0.8 activity threshold on wild-type *E. coli* (screened at 32 µg/ml).^11,16^

### Enrichment factors improve with pre-training across multiple representations

We optimized the virtual screening protocol for antibacterial identification by evaluating the enrichment factor (EF) obtained with different model classes, losses, and transfer-learning regimes. All models in this section were trained on Stokes and evaluated on COADD (molecules from COADD that were structural analogs of those in Stokes were removed).

First, we compared two model classes: a tree-based model on hand-engineered molecular features (XGBoost on Morgan fingerprints) and a DGNN (AttentiveFP) trained with and without transfer learning. Performing this comparison was important because classical models on fingerprints can be highly performant relative to deep neural networks. Figure 3 shows the results of our benchmarking experiment. The poorest level of enrichment was achieved by AttentiveFP without pre-training, thus confirming our hypothesis that the antibacterial data in the training datasets was too small to train DGNNs from scratch; XGBoost on molecular fingerprints achieved fair enrichment, with 2048 bit fingerprints being the most successful. AttentiveFP obtained the highest level of enrichment with pre-training; pre-training was better than training from scratch for all pre-training datasets, and a pre-trained AttentiveFP model was better than XGBoost on fingerprints for almost all pre-training datasets. These results supported our proposed transfer-learning approach for virtual screening.

**Figure 3.**
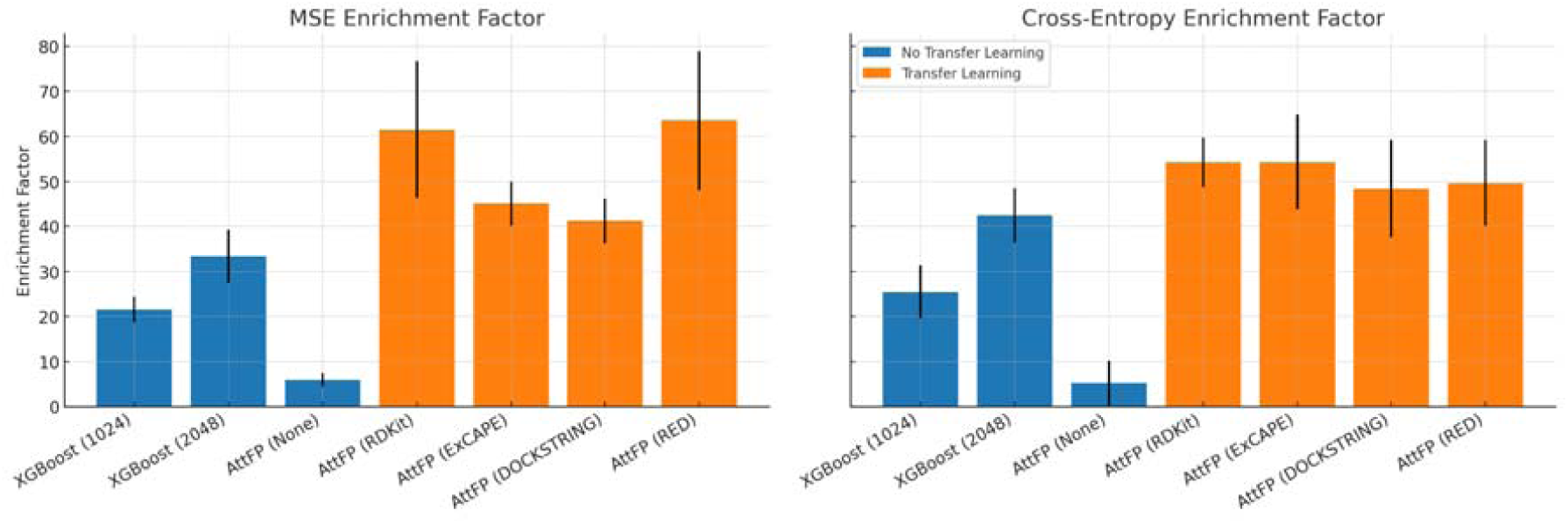
(a) Enrichment factor MSE and (b) Cross-entropy from tree-based models trained on hand-engineered molecular features (XGBoost on Morgan fingerprints) compared to a DGNN (AttentiveFP) trained with and without transfer learning. (Further data in Supplementary Table S1)

Second, we benchmarked different training loss functions. The inhibition values in Stokes and COADD were real numbers ranging primarily between 0 and 1. Prediction of antibacterial activity could, therefore, be framed as regression of raw values, with a mean squared error (MSE) loss, or as classification of binarized values, with a cross-entropy loss. In addition, we tried a custom inhibition loss (IL) that we designed specifically for inhibition since both regression and classification presented disadvantages for predicting inhibition values. This loss was derived from a hard binary classification loss, which was modified by swapping its complex step functions with soft sigmoid-like functions and which was augmented with hyperparameters that controlled the relative weight of false positives and false negatives. Again, we found that pre-training with any of the three losses was superior to training from scratch (Supplementary Table S2). However, we did not observe a significant difference between the three losses, with EFs, which overlapped considerably in terms of standard deviation. Therefore, for simplicity, we decided to frame the prediction of inhibition values as regression with MSE loss.

Finally, we optimized the hyperparameters related to the transfer-learning protocol: the choice of a pre-training dataset, the number of pre-training epochs, and the number of fine-tuning epochs (EF obtained are shown in Supplementary Figure S1). To increase the robustness of our screening protocol, we trained an ensemble of 6 final models for virtual screening. To improve the diversity of models in the ensemble, we selected the two highest-performing hyperparameter combinations and trained three AttentiveFP models with each combination, starting with different random initializations. Since overparameterized neural models are data-intensive, random initialization performs better than bootstrapping the training data when creating ensembles of deep neural models.^46^ The best EF was achieved by pre-training on ExCAPE for 20 epochs and fine-tuning for 500 epochs and by pre-training on RDKit for 10 epochs and fine-tuning for 1000 epochs.

Overall, physicochemical properties demonstrate superior transfer learning effectiveness, with RDKit achieving the highest enrichment factor of 75.2 (10 pre-training epochs, 1000 fine-tuning epochs) compared to binding affinity predictions (ExCAPE: maximum 79.6) and docking scores (DOCKSTRING: maximum 55.6) in the test set (Supplementary Figure S1). While ExCAPE shows the single highest peak performance, RDKit demonstrates more consistent high performance across multiple training configurations (64.3-75.2 range), indicating greater robustness and reliability for transfer learning applications. This could be due to the biological relevance of fundamental molecular descriptors (lipophilicity, hydrogen bonding capacity, molecular size) that influence activity across diverse targets, unlike docking scores which encode highly target-specific geometric complementarity and show the most limited performance ceiling. These results also establish a hierarchical molecular representation framework where physicochemical properties provide the most transferable foundation, binding affinity offers intermediate specificity, and docking scores represent highly specialized but perhaps less transferable features.

### Virtual screening of ultra-large libraries to select structurally novel compounds for experimental testing

Our virtual screening protocol attempted to prioritize the top predictions by the ensemble of DGNNs while maximizing chemical diversity in the final selection to increase the chances of finding hits. First, we predicted the antibacterial activity of every compound in ChemDiv and Enamine. Most predictions were negative, with just a few tens of compounds displaying very high predicted inhibition above 0.8 in each library (Figure 4, a and b). Unsurprisingly, all molecules with very high predicted activity were close analogs of compounds in the training sets Stokes and COADD (Figure 4, c and d), which suggested that our models only predicted very high activity when the candidate molecules belonged to known antibiotic classes. To find antibacterials with novel scaffolds, we expanded our selection to compounds with moderate predicted activity, including all compounds predicted to inhibit growth with values >0.5. Structural novelty was achieved by removing candidates with a Tanimoto similarity to the training set higher than 0.8 (with the exception of a polyketide, a natural product, which was retained as a test compound with known antibiotic properties). Finally, this yielded 151 compounds in ChemDiv and over 10,000 in Enamine, with the caveat that inherent redundancy in combinatorial libraries in Enamine could be responsible for multiple hits.

**Figure 4:**
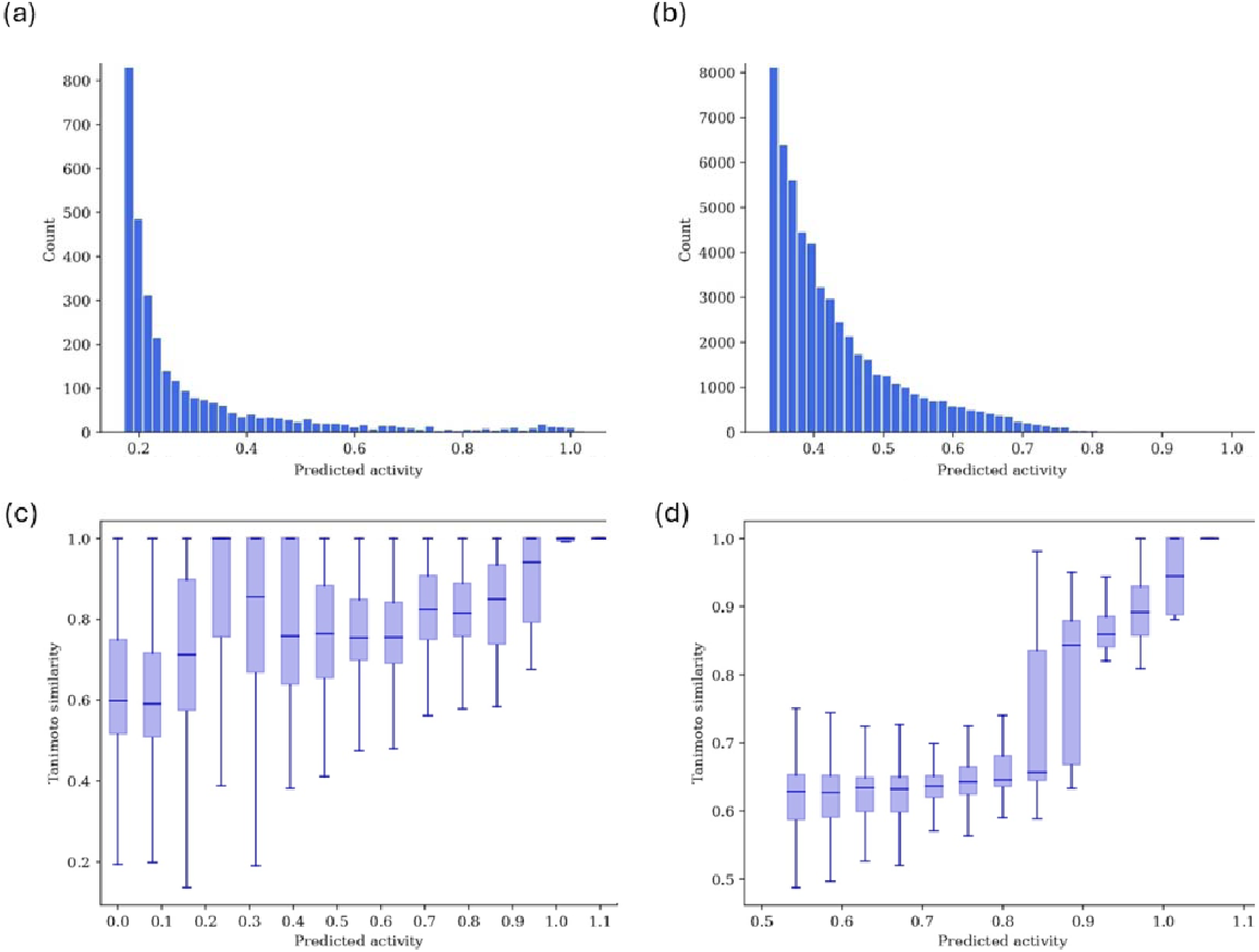
Highest predictions by the ensemble for molecules in (a) ChemDiv and (b) Enamine. Tanimoto’s similarities of the molecules in (c) ChemDiv and (d) Enamine to their closest neighbor in the Stokes and COADD training set are binned by their predicted antibacterial activity. Due to the large size of the Enamine dataset, (d) only shows molecules with predicted inhibition above 0.5. Note that predicted activity outside the [0,1] range were possible: values below 0 suggested that the molecule promoted bacterial growth, while values slightly above 1 could result from experimental variability or error. Count refers to the number of compounds.

To reduce the number of selected compounds while maintaining chemical diversity, we clustered the Enamine pre-selection using molecular fingerprints and selected top-ranking compounds from each cluster to avoid redundancy (as described in *Selection Protocol* in *Methods*) —particularly among overlapping quinolone amides. For both ChemDiv and Enamine, we further grouped compounds into known antibiotic classes or those featuring recurrent functional groups (e.g., nitro moieties), selecting top candidates from each group. The final selection comprised 54 ChemDiv and 140 Enamine compounds, of which 53 and 103, respectively, were available for experimental testing (Table 3), totaling 156 compounds. This included one known polyketide antibiotic retained as a positive control.

**Table 3:**
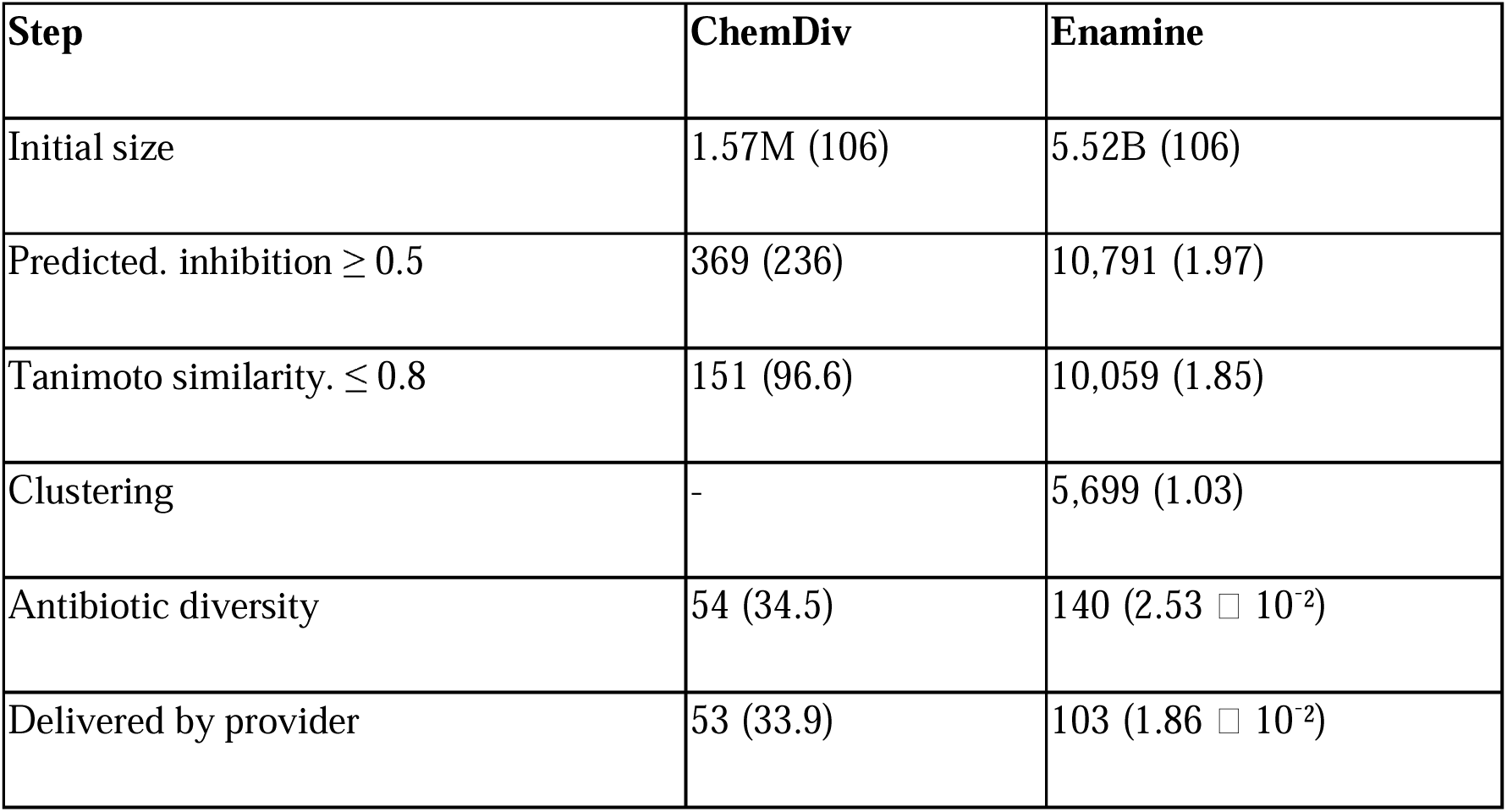
After each selection step, total count (and count per million).

### Visualization of Learned Representations Reveals Antibiotic Class Clustering

To assess whether AttentiveFP models learned meaningful representations for antibiotic classification, we visualized the distribution of compound embeddings in the learned representation space. Embeddings were extracted from one of the models in the ensemble for all compounds in the training datasets Stokes and COADD and in the library ChemDiv. These embeddings, computed by aggregating atom-level representations, allowed us to examine clustering patterns associated with antibiotic classes. Then, we reduced the dimensionality of the embeddings to two dimensions with UMAP. Figure 5 shows the resulting arrangement, highlighting the molecules considered active as per the 0.8 inhibition threshold (measured or predicted), and showing some of the active structures. We observed that clusters grouped similar antibiotic molecules and functional groups, which suggested that our models were learning about the underlying antibiotic classes. However, not all similar antibiotics and functional groups were assigned to the same cluster; for example, quinolones and polyketides were split between two clusters.

**Figure 5:**
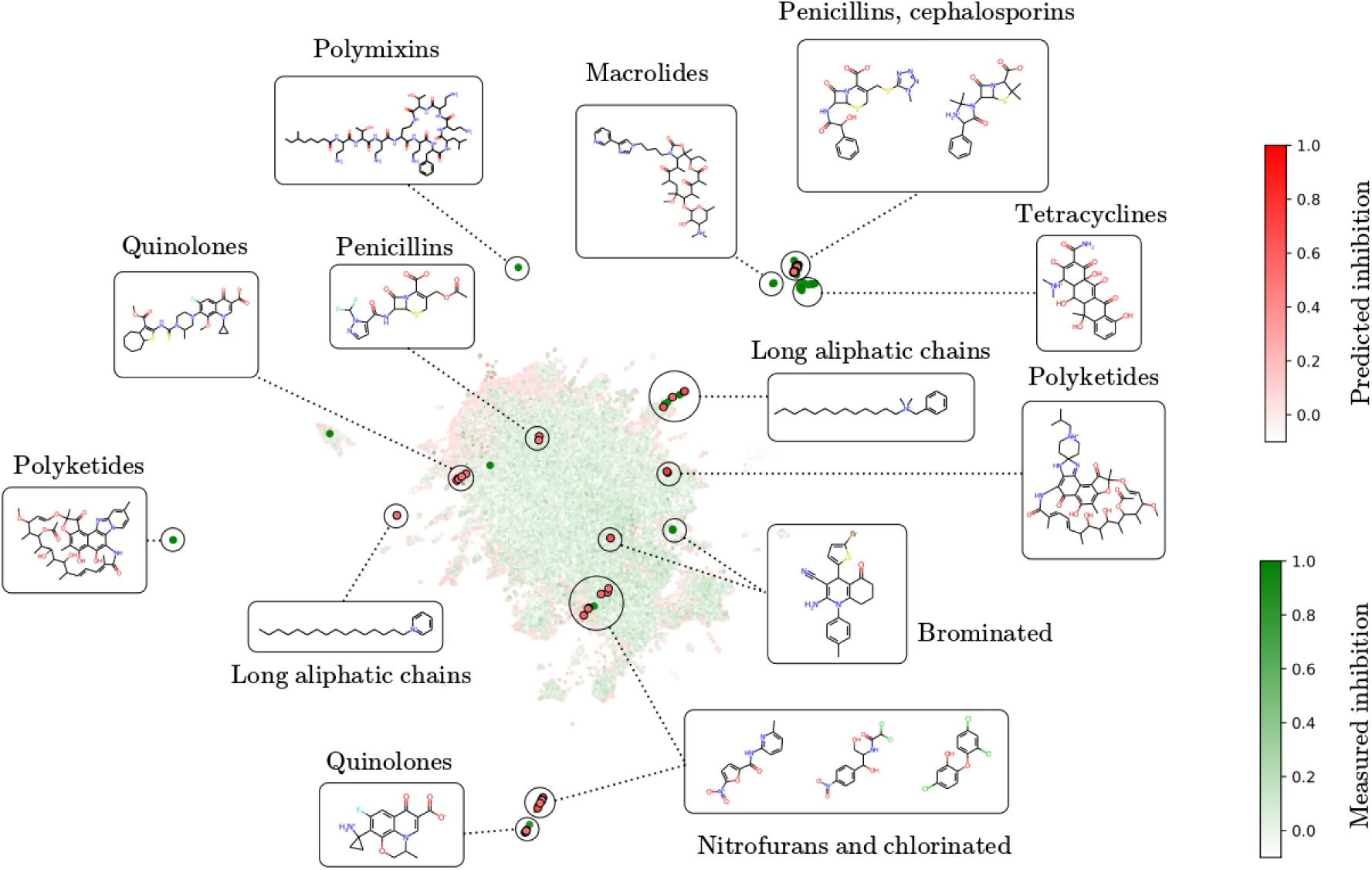
UMAP-reduced visualization of the AttentiveFP embeddings of Stokes, COADD, and ChemDiv. Measured inhibition values from Stokes and COADD are depicted in green, and predicted inhibition from ChemDiv is indicated in red. Molecules with measured or predicted inhibition above 0.8 are highlighted with larger points. Frames show structures representative of each cluster.

### Broad-spectrum testing identifies non-cytotoxic active compounds against gram-positive and gram-negative pathogens

We next tested the minimum inhibitory concentration (MIC) and minimum bactericidal concentration (MBC) of the 156 compounds selected in the previous section against the wild-type strain *E. coli* ATCC 29522. A high level of enrichment for antibacterial activity was observed in the selected subset, with roughly two-thirds of ChemDiv compounds and half of Enamine compounds reaching the MIC hit threshold of 64 µg/ml (Figure 6, Table 4), with a total validation rate of ∼54%. We compared candidates’ MIC and MBC values with those of ampicillin, a known antibiotic used as a control. In our experiment, ampicillin yielded MIC values of 4-8 µg/ml and MBC values of 8-16 µg/ml. Our selection included 10 compounds with better MIC values than ampicillin and 11 with better MBC values (Table 4). The MIC distribution was also observed to have a lower concentration range than the MBC concentration distribution. This was expected because inhibiting growth, as measured by MIC values, is less challenging than killing bacterial cells, as measured by MBC values.

**Figure 6.**
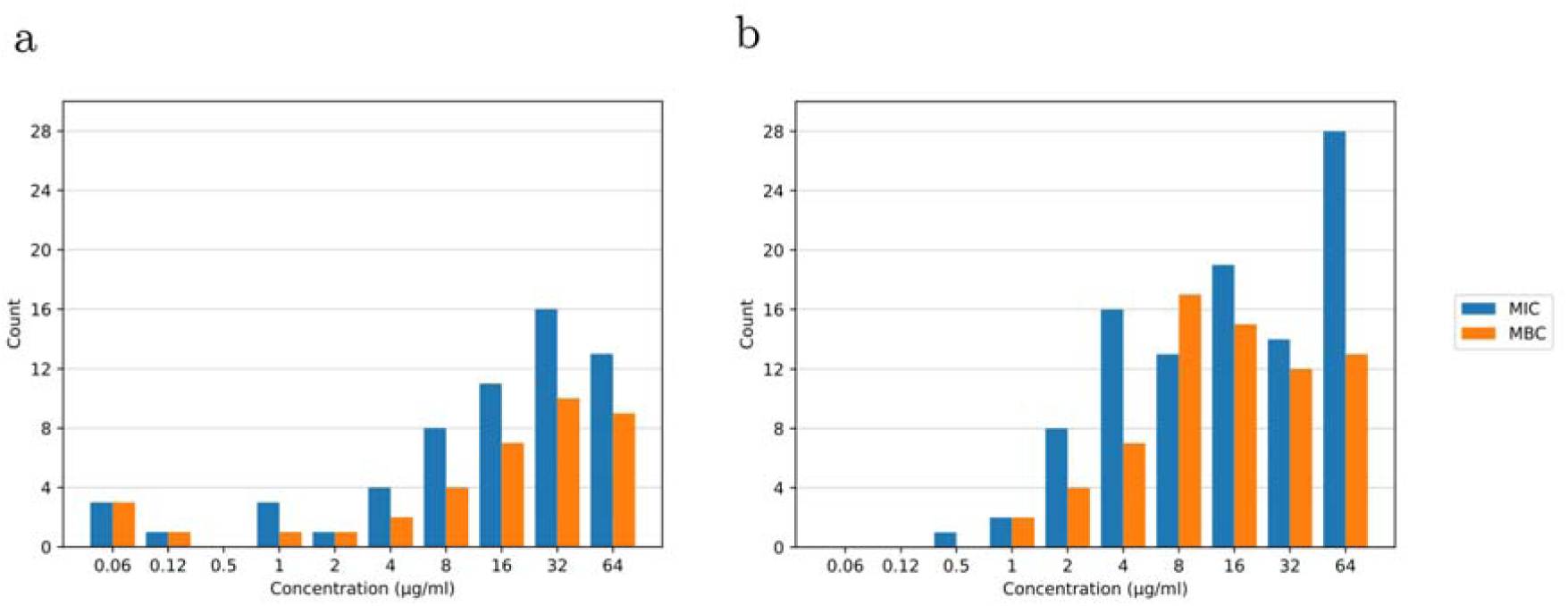
Distribution of MIC and MBC of the (a) 53 molecules selected from ChemDiv and (b) the 103 molecules selected from Enamine; shown only compounds with MIC values of ≤64 µg/ml.

**Table 4.**
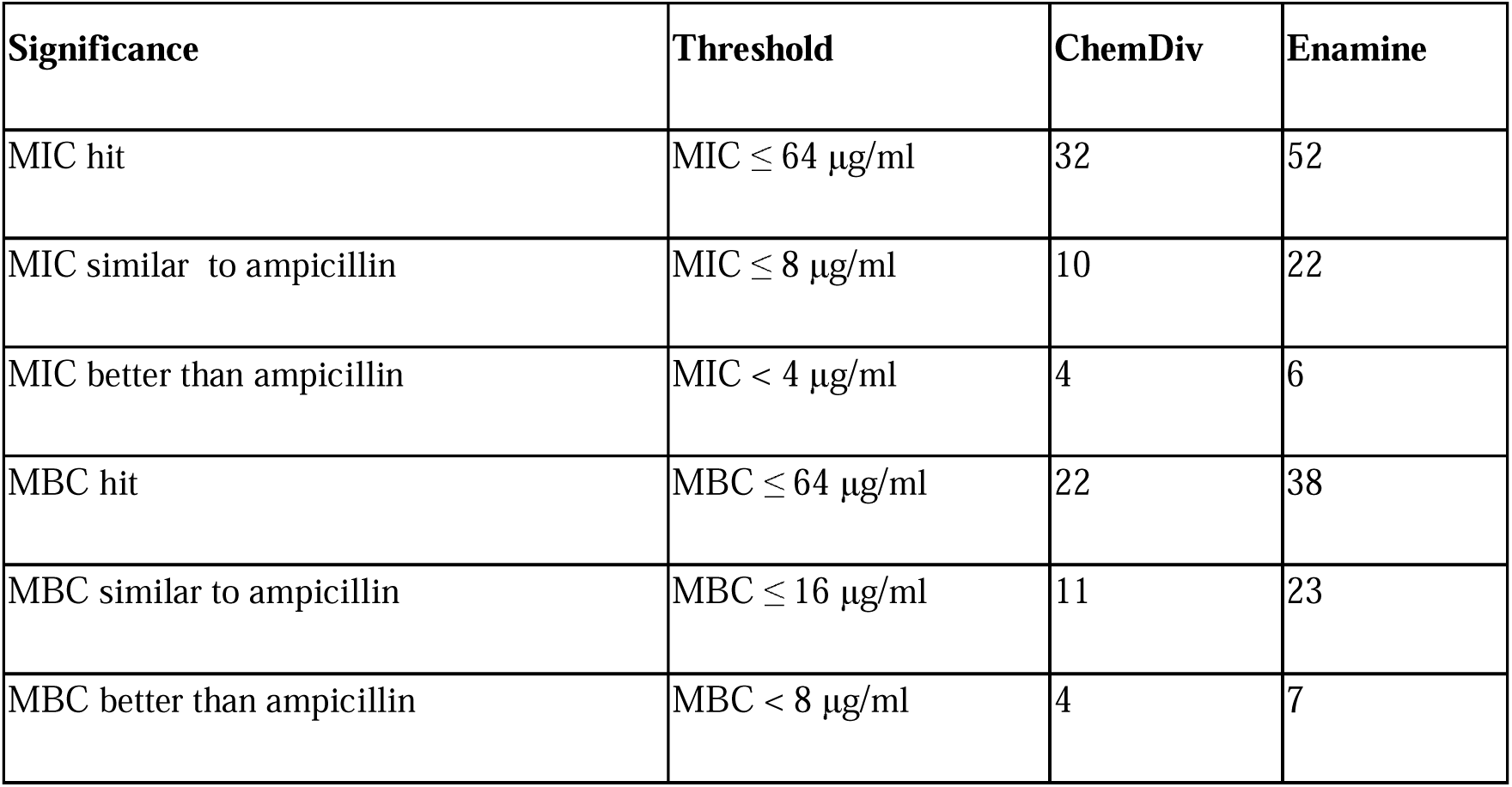
Hits in the 156-molecule selection from ChemDiv and Enamine. Hits are defined as compounds with MICvalues of ≤ 64 μg/ml against *E. coli*. “Similar to ampicillin” is defined as within two-fold of the observed MIC value for ampicillin, or more active than ampicillin.

We next evaluated whether a subset of compounds with activity against *E. col*i had activity against other bacterial species. Though broad-spectrum antibiotics are highly valuable, it is much more straightforward to train models to predict compound activity against a single bacterial strain vs. multiple species. However, given the shared homology of some essential targets and the large proportion of broad-spectrum antibiotics used to train the dataset, it is reasonable that a subset of compounds identified using this model may have broad-spectrum activity.

Due to the high prevalence of quinolone derivatives and nitrofurans in the validated compound set, structural diversity was considered in addition to MIC in the selection of molecules to test for broad-spectrum activity. Compounds were grouped by scaffold or functional group, such as nitrofurans or halogen-containing molecules, and examples from each structural category are shown in Figure 5. A subset of compounds was selected from each category with an MIC requirement of ≤16 µg/ml against *E. coli* (optimising potency, availability and cost among other factors). 18 compounds were evaluated for broad-spectrum activity against the following panel of strains, including three ESKAPE pathogens: a uropathogenic strain of *E. coli* (ECU) 13400, *P. mirabilis* (PM) 432002, *K. pneumoniae* (KP) 13442, *A. baumannii* (AB) 19606 and *S. aureus* (SA) 29213 (for detailed results see Supplementary Table S3).

### Chemical analysis of identified novel antibacterial compounds

Figure 8 shows seven representative examples from the 18 molecules tested, together with their MIC and MBC measurements, the closest molecule in the training set according to the AttentiveFP embeddings, and the most active molecule among the 10 closest molecules in the training set. The top row shows the test compound polyketide with known antibiotic properties which was included in the virtual screening process used as a positive control to test if the workflow selected for active compounds. The bottom row shows the structure of the known antibiotic ampicillin and its MIC and MBC measurements on *E. coli* and *S. aureus*. 16 out of the 18 compounds exhibited better MIC values than ampicillin against either *E. coli* or *S. aureus* (6 out of the 7 in Figure 8), indicating that our protocol was effective at identifying compounds with potent broad-spectrum antibacterial activity. Finally, it is worth noting that the compounds shown here are hits taken directly from the screening library and could potentially be improved through a medicinal chemistry campaign. While some of the hits are simple substitutions of previous antibacterials (e.g., the chlorinated compound 6 in Figure 8), others represent significant deviations from the training set (e.g., compound 5 or compound 7 in Figure 8). As such, they could be compounds that work through a unique mechanism of action compared to current antibiotics, therefore avoiding existing resistance mechanisms.

**Figure 8.**
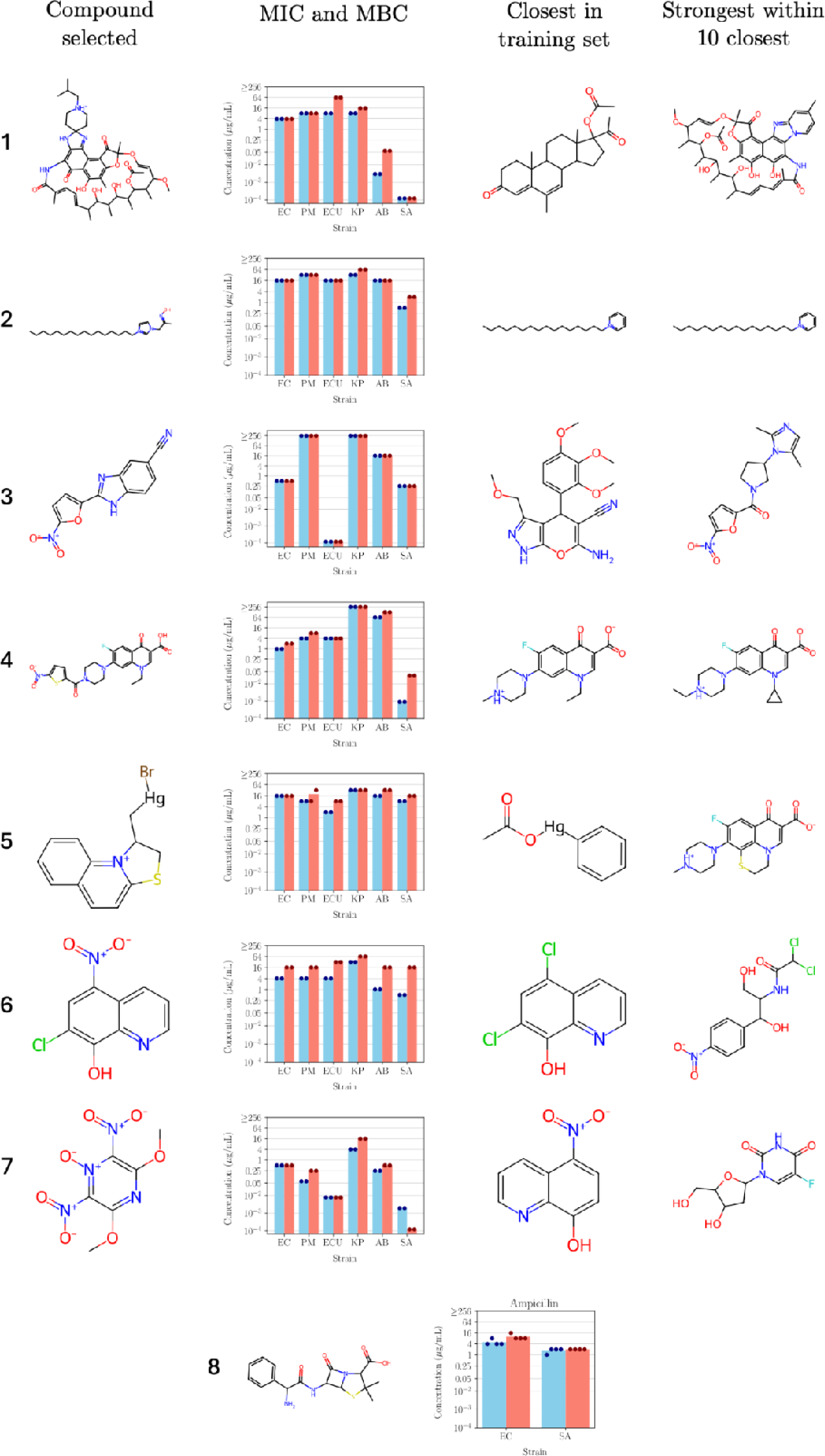
Sample of seven molecules tested in the broad-spectrum validation. The top row shows a polyketide, a natural product that is a known antibiotic used as a positive control included in the virtual screening process to test if the workflow selected for active compounds. The bottom row shows ampicillin, a known antibiotic, also commonly used as the positive control in MIC experiments. Blue bars indicate mean MIC and red bars indicate mean MBC. Dots at the top of the bar indicate the individual measurements from different biological replicates. Distance to the closest compounds in the training set was calculated in the embedding space produced by Attentive FP. For details of all compounds, see Supplementary Table S3.

### Compounds exhibit minimal HepG2 cytotoxicity and hemolytic effects

To ensure that our model was not primarily identifying compounds with general toxicity, these 18 compounds were tested in a cytotoxicity assay against HepG2 cells and a hemolysis assay against erythrocytes. In the HepG2 assay at a concentration of 64 µg/mL, 2 out of 18 compounds demonstrated significant reductions in both cell viability and hemolytic activity. Additionally, one compound showed significant reductions in cell viability only, while another exhibited hemolytic activity only. 15 compounds showed no hemolytic activity at ≤256 µg/ml (maximum concentration tested). Overall, this workflow identified structurally novel compounds with reasonable wild-type antibacterial activity against both gram-positive and gram-negative bacterial species, including multidrug resistant strains, that show limited cytotoxicity.

### Limitations and Future Work

In this study, we used enrichment factors to optimize the virtual screening protocol for antibacterial identification. Enrichment factors have been the established performance metric in virtual screening for decades, specifically designed for scenarios where the objective is to prioritize a small subset of candidates from ultra-large libraries for experimental validation.^47,48^ In the context of antibacterial discovery against ESKAPE pathogens, where sub-micromolar inhibitors are extremely rare, enrichment factors provide the most relevant measure of model performance. We refrain from using general machine learning evaluation metrics such as F1-score, which, while useful in balanced classification tasks, are not well-suited to virtual screening applications. F1-score aggregates precision and recall while accounting for false positives and false negatives. However, in virtual screening for active compound discovery, we operate in an “abundance of discovery” setting where chemical space is vast and screening libraries contain millions of potential compounds. In this context, what matters is discovering compounds that possess the desired biological activity; we do not need to concern ourselves with correctly classifying the enormous number of inactive compounds we choose not to pursue. The drug discovery paradigm fundamentally differs from typical classification problems: false positives are readily identified and filtered out through downstream experimental validation, while false negatives represent missed opportunities for discovering valuable active compounds.^8^ Enrichment factors directly reflect this practical reality by measuring how effectively a model concentrates active compounds in the top-ranked subset relative to random selection, which aligns precisely with the experimental prioritization process. This makes enrichment factors not only more actionable and interpretable for real-world drug discovery applications, but also the historically appropriate metric for this specific field. Our study focuses on virtual screening to discover diverse antibacterial hits from large chemical libraries, rather than serving as a benchmarking exercise for general predictive modeling.

Another limitation of this study is the exclusion of stereochemical information from molecular representations to ensure dataset consistency, as not all publicly available datasets contained complete stereochemical annotations or stereochemistry annotation is poor and absolute configuration is unknown.^8^ While this standardization step is important for fair comparisons, it may reduce model performance, since stereochemistry plays a key role in determining biological activity (enantiomers, for instance, can have vastly different pharmacological effects, toxicities, and target affinities). That said, prior research has shown that 2D molecular representations can often suffice for bioactivity prediction and may even outperform 3D descriptors, as they reduce noise from irrelevant conformational variability and benefit from analogue bias in benchmarks that favor 2D structural similarity over complex 3D features.^49,50^

Our results also highlight the limitations of virtual screening based on existing chemical libraries. Our DGNN ensemble primarily identified compounds structurally related to known antibiotics, such as quinolones, cephalosporins, and penicillins, consistent with prior annotations in the Stokes and COADD training sets. As shown in Figure 5, clusters in the AttentiveFP representation space are aligned with known antibacterial classes. While this validation confirms the model’s learned representation is meaningful, it also implies the screen is biased toward chemical space already covered by known active compounds.

We believe the primary driver was the limited diversity in the fine-tuning data, which overwhelmingly consist of known antibiotic scaffolds. The custom loss function was designed to optimize classification performance, but it did not explicitly penalize structural redundancy or encourage exploration of underrepresented chemical space. Thus, while the loss function may have contributed indirectly to the lack of diversity, the dominant factor appears to be the training data’s chemical composition. Many top Enamine hits were quinolone amides, and although they were active, their structural redundancy necessitated clustering and the selection of representatives. In contrast, ChemDiv yielded more chemically diverse hits, possibly due to its inclusion of natural-product-like scaffolds, despite being ∼3,500 times smaller than Enamine. The inherent redundancy in combinatorial libraries, such as Enamine, may limit their utility unless this diversity is accounted for.

Some hits did deviate substantially from the training space (Figure 6), and these are arguably the most valuable. Finding novel analogs in a fast, automated way with virtual screening could in itself be useful; for example, norfloxacin and ciprofloxacin are both approved for clinical use as antibiotics even though they differ minimally in a single substituent (the former has an ethyl group where the latter has a cyclopropyl). The most valuable hits, however, are those that differ significantly from known antibacterials. Structurally novel antibacterials are more likely to evade known resistance mechanisms, thereby opening up opportunities for new structure-activity relationship (SAR) exploration. They may represent samples from relatively unexplored regions of chemical space. However, our current models only predicted high inhibition for compounds somewhat related to known antibacterials (Figure 5, c and d), which is a limitation shared by many ligand-based virtual screening approaches. To identify relatively novel hits, we needed to consider all candidates with predicted inhibition values higher than 0.5. Other library candidates with high activity but distant from the training set were likely missed in our virtual screening.

The limited structural diversity of known antimicrobials constrains machine learning approaches. Yet, as described by Tommasi et al.^51^, identifying compounds with legitimate activity against wild-type gram-negative bacteria is exceedingly difficult. For example, AstraZeneca screened millions of compounds but was unable to identify any tractable hits against Gram-negative bacteria.^51^

In this work, we did not filter molecules by predicted toxicity or accumulation, and yet our model seems to identify active molecules that are not generally cytotoxic and have some broad-spectrum activity. Although our ensemble approach demonstrated success in the low-data regime for antibacterial prediction, extending this methodology to build reliable predictive models for toxicity profiles would face additional challenges. The heterogeneity in experimental conditions, cell lines, and assay protocols across different toxicity datasets introduces noise.^8^ Furthermore, the integration of multiple toxicity endpoints (cytotoxicity, hemolytic activity, organ-specific toxicity) would require careful consideration of endpoint relationships and potential conflicts between different safety profiles. In the future, a promising direction would be to utilize high-quality data on toxicity and gram-negative accumulation to refine our virtual screening algorithm.

## Conclusions

We developed a virtual screening protocol using ensembles of pre-trained DGNNs to identify antibacterials in large chemical libraries. A key challenge was the scarcity of annotated active compounds—only a few hundred in public datasets—making it difficult for DGNNs trained from scratch to generalize effectively. In this data-sparse regime, classical QSAR models, such as XGBoost, applied to molecular fingerprints, outperformed naïve DGNNs. However, our use of transfer learning dramatically improved model performance. Pre-training on unrelated tasks with hundreds of thousands of molecules enabled the model to learn broadly useful molecular representations. This allowed fine-tuning on small antibacterial datasets to be effective, outperforming XGBoost and DGNNs trained from scratch. While pre-training strategies have been widely successful in other domains (e.g., protein language models, GPT-style architectures^15,52^), their value in drug discovery, particularly for antibacterials, has been more debated. Compared to Stokes et al.^11^ who applied a graph neural network to screen ZINC15, our approach was conceptually more straightforward and achieved high experimental hit rates despite screening a larger number of compounds.

Overall, we demonstrate that transfer learning with deep graph neural networks significantly enhances virtual screening performance in the data-sparse regime of antibacterial discovery. By pre-training on large, general molecular datasets and fine-tuning on limited *E. coli* data, our AttentiveFP ensemble achieved high enrichment factors and identified structurally novel, sub-micromolar compounds active against Gram-positive and Gram-negative ESKAPE pathogens. Experimental validation confirmed a 54% hit rate, with broad-spectrum efficacy and minimal cytotoxicity. The open-source models and scalable workflow developed in this study demonstrates that deep learning models for antibacterial screening can be effectively trained using transfer learning, even when the amount of antibacterial data is limited and the pre-training features are unrelated to antibacterial activity.

## Availability of data and materials

The source code and datasets are available from https://github.com/mgarort/dockbiotic.

## Contributions

M.G. and S.B. defined the project. M.G. designed the model, wrote the code, and ran the experiments. M.G., S.S., and E.G. designed the figures and the experiments. M.G., S.S., and S.B wrote the main manuscript text. C.R., A.B., and S.B provided supervision during the project, with S.B. acting as the primary supervisor. All authors reviewed the manuscript.

## Ethics declarations: Competing interests

The authors have nothing to declare.

## Supplementary Information

### Supplementary Information S1. Inhibition loss function

We developed a custom loss function, called the inhibition loss *l_i_*(IL), which takes as input values in the range [0,1]. The IL was designed to address the limitations of standard regression or classification losses.

The IL is based on a hard binary classification loss (denoted as *l_h_*), which imposes a unit penalty for each false positive or false negative. The hard loss *l_h_* can be written as:

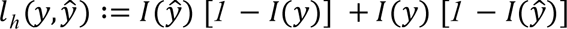

where the first term penalizes false positives and the second term penalizes false negatives. Here, *y* ∈ {0,1} are the true values, 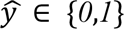 are the predicted values, and *I*(·) is the indicator function:

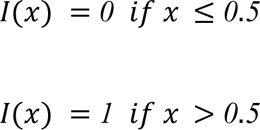

This hard loss *l_h_* can be approximated by a soft loss *l_s_* by replacing the step function witha sigmoid squashing function 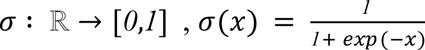

Thus, the soft loss s is:

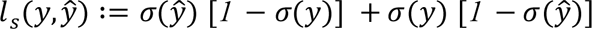

While the sigmoid function σ has its midpoint at *x* = *0, g*(*0*)= *0.5*, this is not always appropriate for all classification tasks. In particular, in our work, we adopted a binarization threshold of 0.8 for antibacterial activity, as suggested by Stokes et al.^11^ To achieve this, we modified the squashing function to ensure that its midpoint occurs at *x* = *0.8*. This modification is represented by:

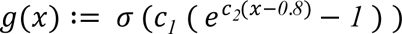

where *c*_1_ controls the range of the squashing function (ensuring *g*(0) ≈ 0 and *g*(1) ≈ 1), and *c*_2_ adjusts the steepness of the slope at *g*(0)= *0.5*. In our experiments, we set *c*_1_ = 4 and *c*_2_ = 3.28.

The final inhibition loss (*l_i_*) is defined as:

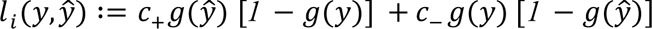

where *c*_+_ and *c*_−_ are coefficients that scale the penalties for false positives and false negatives, respectively. These coefficients were treated as hyperparameters and optimized during training, with values *c*_+_ ∈ [0,1], *c*_−_ ∈ [0,1], and *c*_+_ + *c*_−_ = 1.

## Supplementary Figures

**Supplementary Figure S1.**
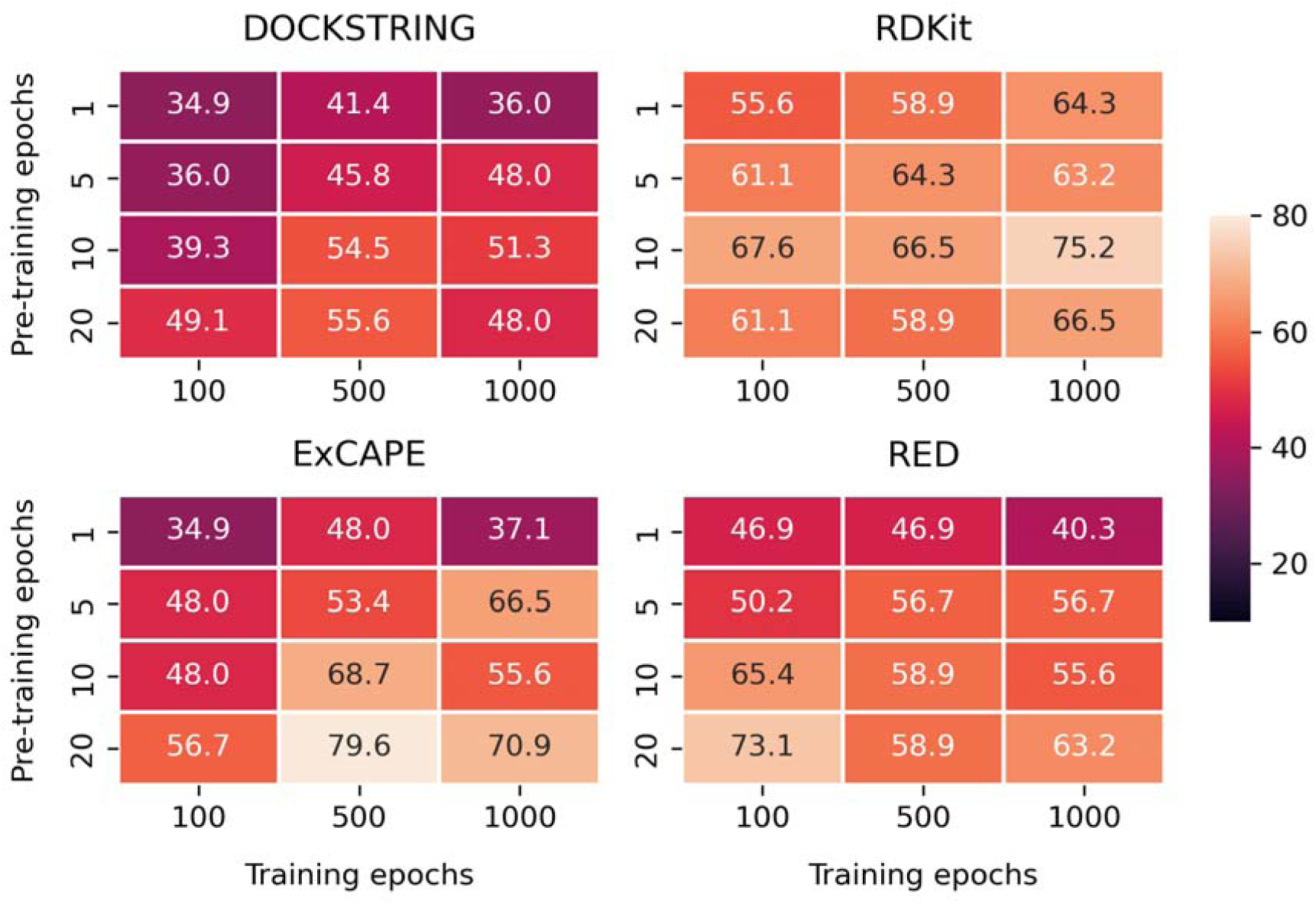
Enrichment Factors obtained in the test set from the various models in the process of optimizing the hyperparameters related to the transfer-learning protocol: the choice of a pre-training dataset, the number of pre-training epochs, and the number of fine-tuning epochs.

## References

1 Antimicrobial Resistance Collaborators, Lancet, 2022, 399, 629–655.

2 M. E. A. de Kraker, A. J. Stewardson and S. Harbarth, PLoS Med., 2016, 13, e1002184.

3 M. K. Gugger and P. J. Hergenrother, Nature, 2024, 625, 451–452.

4 K. Lewis, Nat. Rev. Drug Discov., 2013, 12, 371–387.

5 E. D. Brown and G. D. Wright, Nature, 2016, 529, 336–343.

6 M. A. Cooper and D. Shlaes, Nature, 2011, 472, 32.

7 M. A. T. Blaskovich, J. Zuegg, A. G. Elliott and M. A. Cooper, ACS Infect. Dis., 2015, 1, 285–287.

8 10.1021/acs.chemrestox.5c00033, (accessed 20 April 2025).

9 A. Liu, S. Seal, H. Yang and A. Bender, SLAS Discov., 2023, 28, 53–64.

10 E. Lawrence, A. El-Shazly, S. Seal, C. K. Joshi, P. Liò, S. Singh, A. Bender, P. Sormanni and M. Greenig, arXiv [cs. AI].

11 J. M. Stokes, K. Yang, K. Swanson, W. Jin, A. Cubillos-Ruiz, N. M. Donghia, C. R. MacNair, S. French, L. A. Carfrae, Z. Bloom-Ackermann, V. M. Tran, A. Chiappino-Pepe, A. H. Badran, I. W. Andrews, E. J. Chory, G. M. Church, E. D. Brown, T. S. Jaakkola, R. Barzilay and J. J. Collins, Cell, 2020, 181, 475–483.

12 F. Wong, E. J. Zheng, J. A. Valeri, N. M. Donghia, M. N. Anahtar, S. Omori, A. Li, A. Cubillos-Ruiz, A. Krishnan, W. Jin, A. L. Manson, J. Friedrichs, R. Helbig, B. Hajian, D. K. Fiejtek, F. F. Wagner, H. H. Soutter, A. M. Earl, J. M. Stokes, L. D. Renner and J. J. Collins, Nature, DOI:10.1038/s41586-023-06887-8.

13 A. Krizhevsky, I. Sutskever and G. E. Hinton, Commun. ACM, 2017, 60, 84–90.

14 C. Saharia, W. Chan, S. Saxena, L. Li, J. Whang, E. Denton, S. K. S. Ghasemipour, B. K. Ayan, S. S. Mahdavi, R. G. Lopes, T. Salimans, J. Ho, D. J. Fleet and M. Norouzi, arXiv [cs.CV], 2022.

15 OpenAI, J. Achiam, S. Adler, S. Agarwal, L. Ahmad, I. Akkaya, F. L. Aleman, D. Almeida, J. Altenschmidt, S. Altman, S. Anadkat, R. Avila, I. Babuschkin, S. Balaji, V. Balcom, P. Baltescu, H. Bao, M. Bavarian, J. Belgum, I. Bello, J. Berdine, G. Bernadett-Shapiro, C. Berner, L. Bogdonoff, O. Boiko, M. Boyd, A.-L. Brakman, G. Brockman, T. Brooks, M. Brundage, K. Button, T. Cai, R. Campbell, A. Cann, B. Carey, C. Carlson, R. Carmichael, B. Chan, C. Chang, F. Chantzis, D. Chen, S. Chen, R. Chen, J. Chen, M. Chen, B. Chess, C. Cho, C. Chu, H. W. Chung, D. Cummings, J. Currier, Y. Dai, C. Decareaux, T. Degry, N. Deutsch, D. Deville, A. Dhar, D. Dohan, S. Dowling, S. Dunning, A. Ecoffet, A. Eleti, T. Eloundou, D. Farhi, L. Fedus, N. Felix, S. P. Fishman, J. Forte, I. Fulford, L. Gao, E. Georges, C. Gibson, V. Goel, T. Gogineni, G. Goh, R. Gontijo-Lopes, J. Gordon, M. Grafstein, S. Gray, R. Greene, J. Gross, S. S. Gu, Y. Guo, C. Hallacy, J. Han, J. Harris, Y. He, M. Heaton, J. Heidecke, C. Hesse, A. Hickey, W. Hickey, P. Hoeschele, B. Houghton, K. Hsu, S. Hu, X. Hu, J. Huizinga, S. Jain, S. Jain, J. Jang, A. Jiang, R. Jiang, H. Jin, D. Jin, S. Jomoto, B. Jonn, H. Jun, T. Kaftan, Ł. Kaiser, A. Kamali, I. Kanitscheider, N. S. Keskar, T. Khan, L. Kilpatrick, J. W. Kim, C. Kim, Y. Kim, J. H. Kirchner, J. Kiros, M. Knight, D. Kokotajlo, Ł. Kondraciuk, A. Kondrich, A. Konstantinidis, K. Kosic, G. Krueger, V. Kuo, M. Lampe, I. Lan, T. Lee, J. Leike, J. Leung, D. Levy, C. M. Li, R. Lim, M. Lin, S. Lin, M. Litwin, T. Lopez, R. Lowe, P. Lue, A. Makanju, K. Malfacini, S. Manning, T. Markov, Y. Markovski, B. Martin, K. Mayer, A. Mayne, B. McGrew, S. M. McKinney, C. McLeavey, P. McMillan, J. McNeil, D. Medina, A. Mehta, J. Menick, L. Metz, A. Mishchenko, P. Mishkin, V. Monaco, E. Morikawa, D. Mossing, T. Mu, M. Murati, O. Murk, D. Mély, A. Nair, R. Nakano, R. Nayak, A. Neelakantan, R. Ngo, H. Noh, L. Ouyang, C. O’Keefe, J. Pachocki, A. Paino, J. Palermo, A. Pantuliano, G. Parascandolo, J. Parish, E. Parparita, A. Passos, M. Pavlov, A. Peng, A. Perelman, F. de Avila Belbute Peres, M. Petrov, H. P. de Oliveira Pinto, Michael, Pokorny, M. Pokrass, V. H. Pong, T. Powell, A. Power, B. Power, E. Proehl, R. Puri, A. Radford, J. Rae, A. Ramesh, C. Raymond, F. Real, K. Rimbach, C. Ross, B. Rotsted, H. Roussez, N. Ryder, M. Saltarelli, T. Sanders, R. Santurkar, G. Sastry, H. Schmidt, D. Schnurr, J. Schulman, D. Selsam, K. Sheppard, R. Sherbakov, J. Shieh, S. Shoker, P. Shyam, S. Sidor, E. Sigler, M. Simens, J. Sitkin, K. Slama, I. Sohl, B. Sokolowsky, Y. Song, N. Staudacher, F. P. Such, N. Summers, I. Sutskever, J. Tang, N. Tezak, M. B. Thompson, P. Tillet, A. Tootoonchian, E. Tseng, P. Tuggle, N. Turley, J. Tworek, J. F. C. Uribe, A. Vallone, A. Vijayvergiya, C. Voss, C. Wainwright, J. J. Wang, A. Wang, B. Wang, J. Ward, J. Wei, C. J. Weinmann, A. Welihinda, P. Welinder, J. Weng, L. Weng, M. Wiethoff, D. Willner, C. Winter, S. Wolrich, H. Wong, L. Workman, S. Wu, J. Wu, M. Wu, K. Xiao, T. Xu, S. Yoo, K. Yu, Q. Yuan, W. Zaremba, R. Zellers, C. Zhang, M. Zhang, S. Zhao, T. Zheng, J. Zhuang, W. Zhuk and B. Zoph, arXiv [cs.CL], 2023.

16 M. R. Desselle, R. Neale, K. A. Hansford, J. Zuegg, A. G. Elliott, M. A. Cooper and M. A. Blaskovich, Future Sci. OA, 2017, 3, FSO171.

17 D. Buterez, J. P. Janet, S. J. Kiddle, D. Oglic and P. Lió, Nat. Commun., 2024, 15, 1517.

18 Y. Rong, Y. Bian, T. Xu, W. Xie, Y. Wei, W. Huang and J. Huang, arXiv [q-bio.BM], 2020.

19 F. Zhuang, Z. Qi, K. Duan, D. Xi, Y. Zhu, H. Zhu, H. Xiong and Q. He, arXiv [cs.LG], 2019.

20 X. Li and D. Fourches, J. Cheminform., 2020, 12, 27.

21 E. King-Smith, ChemRxiv, 2023.

22 A. Keto, T. Guo, N. Gönnheimer, X. Zhang, E. H. Krenske and O. Wiest, Digit. Discov., 2025, 4, 1232–1238.

23 N. Noto, R. Kunisada, T. Rohlfs, M. Hayashi, R. Kojima, O. García Mancheño, T. Yanai and S. Saito, Nat. Commun., 2025, 16, 3388.

24 M. Sypetkowski, F. Wenkel, F. Poursafaei, N. Dickson, K. Suri, P. Fradkin and D. Beaini, arXiv [cs.LG], 2024.

25 M. García-Ortegón, G. N. C. Simm, A. J. Tripp, J. M. Hernández-Lobato, A. Bender and S. Bacallado, J. Chem. Inf. Model., 2022, 62, 3486–3502.

26 M. García-Ortegón, S. Seal, C. Rasmussen, A. Bender and S. Bacallado, J. Cheminform., 2024, 16, 115.

27 O. Méndez-Lucio, C. A. Nicolaou and B. Earnshaw, Nat. Commun., 2024, 15, 9431.

28 H. Li, R. Zhang, Y. Min, D. Ma, D. Zhao and J. Zeng, Nat. Commun., 2023, 14, 7568.

29 J. Sun, N. Jeliazkova, V. Chupakhin, J.-F. Golib-Dzib, O. Engkvist, L. Carlsson, J. Wegner, H. Ceulemans, I. Georgiev, V. Jeliazkov, N. Kochev, T. J. Ashby and H. Chen, J. Cheminform., DOI:10.1186/s13321-017-0203-5.

30 G. Landrum, Greg Landrum, 2013, 8, 5281.

31 N. M. O’Boyle, M. Banck, C. A. James, C. Morley, T. Vandermeersch and G. R. Hutchison, J. Cheminform., 2011, 3, 33.

32 T. Chen and C. Guestrin, arXiv [cs.LG], 2016.

33 H. L. Morgan, J. Chem. Doc., 1965, 5, 107–113.

34 D. Rogers and M. Hahn, J. Chem. Inf. Model., 2010, 50, 742–754.

35 Z. Xiong, D. Wang, X. Liu, F. Zhong, X. Wan, X. Li, Z. Li, X. Luo, K. Chen, H. Jiang and M. Zheng, J. Med. Chem., 2020, 63, 8749–8760.

36 B. Ramsundar, K. Leswing, P. Eastman and V. Pande, Deep learning for the life sciences, O’Reilly Media, Sebastopol, CA, 2019.

37 D. P. Kingma and J. Ba, arXiv [cs.LG], 2014.

38 CHEMDIV INC - FULLY INTEGRATED TARGET-TO-CLINIC CONTRACT RESEARCH ORGANIZATION, https://www.chemdiv.com, (accessed 29 December 2024).

39 O. O. Grygorenko, European J. Org. Chem., 2021, 2021, 6474–6477.

40 Chemical Supplier - Enamine, https://enamine.net, (accessed 29 December 2024).

41 REAL Space, https://enamine.net/compound-collections/real-compounds/real-space-navigator, (accessed 29 December 2024).

42 M. Ester, H. Kriegel, J. Sander and X. Xu, KDD, 1996, 226–231.

43 R. Gómez-Bombarelli, J. N. Wei, D. Duvenaud, J. M. Hernández-Lobato, B. Sánchez-Lengeling, D. Sheberla, J. Aguilera-Iparraguirre, T. D. Hirzel, R. P. Adams and A. Aspuru-Guzik, ACS Cent Sci, 2018, 4, 268–276.

44 L. McInnes, J. Healy and J. Melville, arXiv [stat.ML], 2018.

45 L. J. Farrell, R. Lo, J. J. Wanford, A. Jenkins, A. Maxwell and L. J. V. Piddock, J. Antimicrob. Chemother., 2018, 73, 2284–2297.

46 B. Lakshminarayanan, A. Pritzel and C. Blundell, arXiv [stat.ML], 2016.

47 A. Bender and R. C. Glen, J. Chem. Inf. Model., 2005, 45, 1369–1375.

48 S. M. Paul, D. S. Mytelka, C. T. Dunwiddie, C. C. Persinger, B. H. Munos, S. R. Lindborg and A. L. Schacht, Nat. Rev. Drug Discov., 2010, 9, 203–214.

49 K. Gao, D. D. Nguyen, V. Sresht, A. M. Mathiowetz, M. Tu and G.-W. Wei, Phys. Chem. Chem. Phys., 2020, 22, 8373–8390.

50 M. S. Bahia, O. Kaspi, M. Touitou, I. Binayev, S. Dhail, J. Spiegel, N. Khazanov, A. Yosipof and H. Senderowitz, Mol. Inform., 2023, 42, e2200186.

51 R. Tommasi, D. G. Brown, G. K. Walkup, J. I. Manchester and A. A. Miller, Nat. Rev. Drug Discov., 2015, 14, 529–542.

52 E. Nguyen, M. Poli, M. G. Durrant, A. W. Thomas, B. Kang, J. Sullivan, M. Y. Ng, A. Lewis, A. Patel, A. Lou, S. Ermon, S. A. Baccus, T. Hernandez-Boussard, C. Re, P. D. Hsu and B. L. Hie, bioRxiv, 2024.

